# CyclicMPNN: Stable Cyclic Peptide Sequence Generation

**DOI:** 10.64898/2026.01.31.702993

**Authors:** Andrew C. Powers, Yanapat Janthana, Parisa Hosseinzadeh

## Abstract

Cyclic peptides are a promising class of therapeutics due to their attractive drug qualities such as increased structural stability, cell permeability, and resistance to proteolytic degradation. With recent advancements in cyclic peptide backbone generation models like CyclicCAE and RFPeptide, generating cyclic peptide backbones can be done more rapidly compared to traditional algorithm or physics based approaches. However, designing energetically favorable cyclic peptide sequences to fit generated backbones using only canonical amino acids is nontrivial. We fine-tuned the state-of-the-art deep learning model for protein sequence design, ProteinMPNN, using a combination of X-ray crystal structures from the Protein Data Bank and *in silico* generated cyclic peptides. Our approach surpasses ProteinMPNN in cyclic peptide sequence design, producing energetically stable sequences with a higher success rate of folding into the generated cyclic peptide backbones. We show that CyclicMPNN can be used as a motif-inpainting strategy and in *de novo* sequence design tasks. We propose that CyclicMPNN will enable the rapid design of energetically stable cyclic peptide sequences, increasing the success rate of therapeutic cyclic peptide development.

## Introduction

Cyclic peptides are a class of therapeutics that have attracted increasing interest due to their enhanced structural stability [1], cell permeability [2, 3, 4, 5, 6, 7], and resistance to proteolytic degradation [6, 4]. Cyclic peptides can be synthesized chemically using solid-phase peptide synthesis [8, 9], as well as through phage display libraries [10, 11, 12]. These synthesis approaches enable tuning of chemical properties to improve stability, affinity, specificity, and permeability.

Traditionally, computational design of cyclic peptides has been largely limited to physics-based methods such as Rosetta [13, 14, 15, 16] and molecular dynamics (MD) simulations [17, 18]. In these frameworks, sequences can be generated *de novo* or rationally designed, and conformational ensembles of designed cyclic peptides can be sampled in solution, though often at high computational cost. In Rosetta, multiple approaches [19] can be used to generate structural ensembles for a given peptide sequence through backbone sampling based on amino acid Ramachandran preferences. In MD, conformational sampling is commonly performed using enhanced sampling techniques such as bias-exchange metadynamics (BE-META) or replica exchange molecular dynamics (REMD) [20].

Deep learning models developed for protein sequence design, such as the state-of-the-art ProteinMPNN [21], can also be applied to cyclic peptides. Although ProteinMPNN was not explicitly trained on peptides or cyclic peptides, it has been used to generate stable cyclic peptides in combination with iterative design and Rosetta relaxation [22, 23]. However, this workflow typically requires multiple rounds of design and structure refinement. A ProteinMPNN model with weights specialized for cyclic peptides would therefore be desirable to improve efficiency and performance. Several variants of ProteinMPNN have been developed for specific design contexts, including ThermoMPNN [24], SolubleMPNN [25], LigandMPNN [26], and HighMPNN [27], the latter incorporating an additional module to enhance cyclic peptide sequence generation.

Here, we present CyclicMPNN, a fine-tuned version of ProteinMPNN developed specifically for cyclic peptide sequence design. ProteinMPNN was fine-tuned using a mixture of Protein Data Bank (PDB) structures and *in silico* generated cyclic peptide backbones. The *in silico* dataset was created using Generalized Kinematic loop Closure (GenKIC), a loop conformation sampling approach (see Methods), to generate cyclic poly-alanine backbone ensembles. Sequences were assigned to these backbones using ProteinMPNN, and the resulting sequences were folded using a modified AlphaFold model, HighFold [28], which replaces linear positional encodings with relative positional encodings through modification of the model’s offset matrix.

CyclicMPNN shows improved structural reconstruction accuracy, as measured by RMSD, and higher predicted Local Distance Difference Test (pLDDT) scores compared to ProteinMPNN and HighMPNN when designing sequences for *de novo* cyclic peptide backbones, independent of the backbone sampling method. CyclicMPNN designed sequences also exhibit greater sequence diversity and reduced repetition relative to ProteinMPNN. For experimentally determined cyclic peptide structures, CyclicMPNN achieves lower predicted RMSD to X-ray backbones in a single round of sequence design. In motif-inpainting tasks, CyclicMPNN further outperforms ProteinMPNN, producing approximately twice as many stable cyclic peptides containing the desired motif.

We propose that CyclicMPNN represents a state-of-the-art approach for cyclic peptide sequence design and provides a practical tool for high-throughput cyclic peptide engineering, analogous to the impact of ProteinMPNN on protein design. More broadly, our dataset generation and fine-tuning strategy offers a general framework for extending pretrained protein design models to under represented structural regimes and other specialized design tasks.

## Results

### CyclicMPNN can generate novel sequences that recapitulate x-ray crystal structure

We generated in silico data and fine-tuned ProteinMPNN, resulting in an overall improvement in cyclic peptide sequence design performance (Fig. 1A & 1B; details in the Methods section). As an initial test of CyclicMPNN’s ability to sample sequences consistent with experimentally observed cyclic peptides, we evaluated which design method sampled sequences compatible with cyclic peptides whose structures were determined by X-ray crystallography and deposited in the PDB. Four crystal structures (9CDT, 9HGC, 9CDU, 9HGD) from Rettie et al. (2025) were selected as representative test cases. These peptides were designed using a combination of RFPeptide [29], backbone conformation ensemble generation, and ProteinMPNN for sequence design. ProteinMPNN and HighMPNN were used to benchmark CyclicMPNN’s performance. Given that these peptides were originally designed using iterative rounds of ProteinMPNN, we initially expected ProteinMPNN to outperform CyclicMPNN when redesigning sequences for these structures. Predicted structures for sampled sequences were generated using HighFold [28].

**Figure 1:**
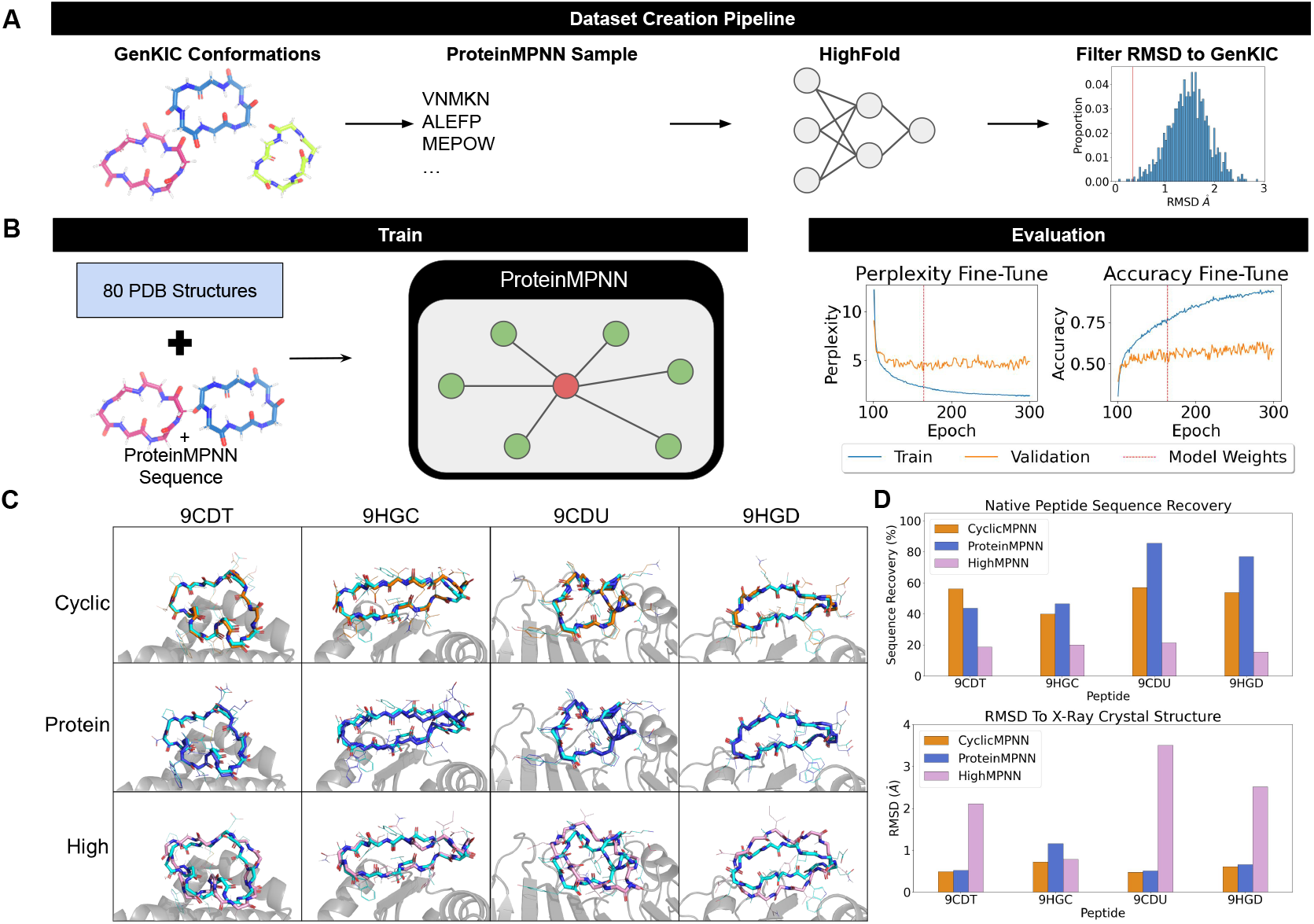
Overview of CyclciMPNN data creation and fine-tuning, with X-ray crystal structure comparison. **A)** Data creation representation where we generate poly-alanine peptide conformations using GenKIC, designed sequences using ProteinMPNN, folded the sequences using HighFold, and then compared to GenKIC structures. **B)** ProteinMPNN fine-tuning methodology where we took our *in silico* generated data and 80 structures from the Protein Data Back. Weights were picked off of lowest validation perplexity loss and before over training. **C)** Four crystal structures (9CDT, 9HGC, 9CDU, 9HGD) were used to benchmark CyclicMPNN (orange), ProteinMPNN (blue), and HighMPNN (plum) sequence generation. Predicted structures to the X-ray crystal structure (teal) are shown aligned. **D)** Top: Overview of how unique our closest RMSD sequence were to the native sequence, Bottom: RMSD comparison of structures against native crystal structures.

When redesigning sequences for the four peptides, CyclicMPNN outperformed ProteinMPNN and HighMPNN in both structural reconstruction accuracy (RMSD) and sequence uniqueness (Fig. 1C & 1D). Notably, ProteinMPNN’s lowest RMSD sequences showed variable sequence recovery relative to the native peptides, with lower recovery for two structures (44% and 46%) and higher recovery for the other two (86% and 77%). ProteinMPNN achieved low backbone RMSDs overall (0.5 Å, 1.2 Å, 0.5 Å, and 0.7 Å), with only one structure exceeding 1 Å. HighMPNN showed the lowest sequence recovery overall (19%, 20%, 21%, and 15%) and higher structural deviations, with RMSDs of 2.0 Å, 0.8 Å, 3.5 Å, and 2.5 Å; only one structure had RMSD *≤* 1 Å. CyclicMPNN exhibited more consistent sequence recovery (50%; 56%, 40%, 57%, and 54%). All CyclicMPNN designed sequences yielded high-accuracy structural reconstructions (RMSD *≤* 1.0 Å; 0.5 Å, 0.7 Å, 0.5 Å, and 0.6 Å).

### CyclicMPNN produces better sequences for *de novo* structure than ProteinMPNN

ProteinMPNN is commonly used as a sequence design method for newly generated or diffused backbones. To evaluate CyclicMPNN’s sequence design performance on *de novo* structures, we tested both methods on synthetic backbone ensembles. Since multiple machine learning and physics-based approaches exist for generating poly-alanine or poly-glycine backbone conformation ensembles of different sizes, this provides a controlled framework for assessing how effectively CyclicMPNN can generate sequences for a given scaffold.

Cyclic peptide ensembles of size 10,000 of 6, 8, and 10 amino acids were generated using GenKIC (methodology in Methods) with poly-alanine backbones. Using poly-alanine structures ensures that the conformational ensembles reflect dihedral preferences of all L-*α*-amino acids, as CyclicMPNN, ProteinMPNN, and HighMPNN do not design sequences containing non-canonical amino acids.

To determine which method produced more reliable sequences, one sequence was generated for each of the 10,000 backbone structures at each peptide length. Example predicted structures from sequences designed by both methods for the 6-, 8-, and 10-residue GenKIC datasets are shown in Fig. 2A. CyclicMPNN sequences more closely recapitulated the target *de novo* structures compared to ProteinMPNN designed sequences. Predicted structures were generated using HighFold, and pLDDT and RMSD values were computed and compared (Fig. 2B, left column). On average, CyclicMPNN outperformed ProteinMPNN in both RMSD and pLDDT across all sizes. The median RMSDs for 6-, 8-, and 10-mers were 0.97 Å vs. 1.17 Å, 1.13 Å vs. 1.55 Å, and 1.79 Å vs. 1.99 Å for CyclicMPNN and ProteinMPNN, respectively. Median pLDDT values for 6-, 8-, and 10-mers were 80.13 vs. 76.18, 82.00 vs. 74.86, and 71.91 vs. 70.18 for CyclicMPNN and ProteinMPNN, respectively. Across all three lengths, CyclicMPNN designed sequences more frequently adopted the intended backbone conformation. Relative improvements in structural recapitulation for CyclicMPNN over ProteinMPNN were 16.23%, 20.60%, and 9.23% for 6-, 8-, and 10-mers, respectively.

**Figure 2:**
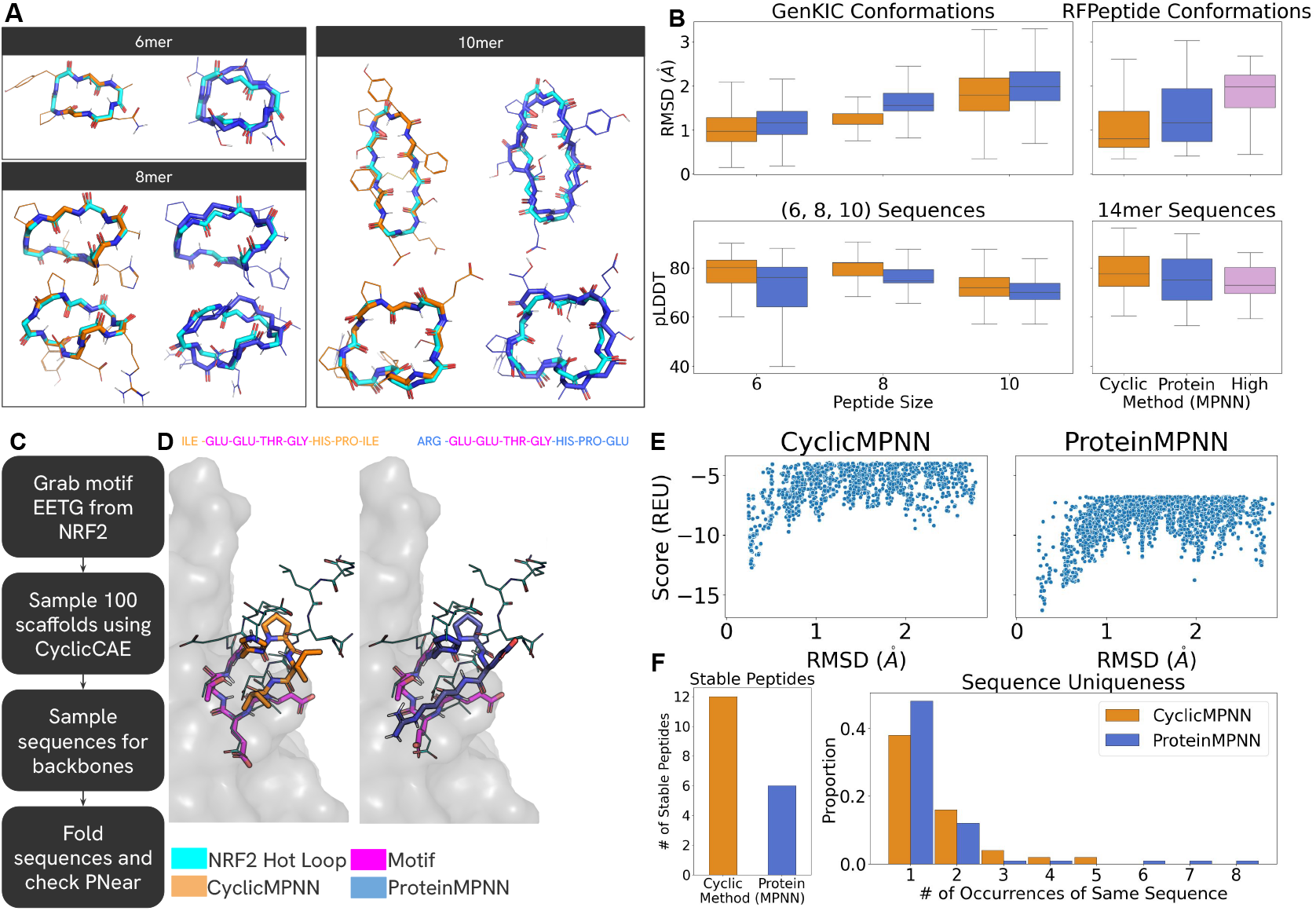
CyclicMPNN outperform ProteinMPNN and HighMPNN for *De Novo* Sequence design tasks and motif-inpainting tasks. **A)** *De novo* conformer structures of size 6, 8, and 10 amino acids generated (teal) with CyclicMPNN (orange) and ProteinMPNN (blue) predicted structures, based on sequences produced, are overlaid. **B)** Left: Boxplot of RMSD & pLDDT metrics from our 10,000 predicted structures, based on the sequences from CyclicMPNN and ProteinMPNN, compared to our GenKIC structures. Right: RMSD & pLDDT of the 100 CyclicMPNN, ProteinMPNN, and HighMPNN predicted structures compared to the generated RFPeptide poly-glycine 14-mers used for design. Workflow for producing motif inpainting structures using the hot loop based off of Nrf2 **D)** Overlaid structural predicitions for CyclicMPNN and ProteinMPNN from `simple_cycpep_predict` overlaid onto the Nrf2 motif. **E)** P_*Near*_ graphs of both CyclicMPNN (left) and ProteinMPNN (right) shown in D. **F)** Left: Number of stable sequences compared between methods. Right: Number of occurrences for each sequence between the two methods when motif-inpainting for Nrf2.

For 6-mers, we additionally selected 100 sequences with RMSD *≤* 0.5 Å relative to the original poly-alanine backbones. *P*_*Near*_ stability was calculated for this subset (Supplementary Fig. S1). CyclicMPNN sequences showed a statistically significantly higher median *P*_*Near*_ value (0.6) compared to ProteinMPNN (0.08).

To evaluate larger cyclic peptides, we used RFPeptide to generate 14-mer backbone conformations (Fig. 2B, right column). CyclicMPNN again achieved lower median RMSD and higher median pLDDT compared to ProteinMPNN and HighMPNN. Median RMSDs for RFPeptide backbones were 0.81 Å, 1.16 Å, and 1.98 Å for CyclicMPNN, ProteinMPNN, and HighMPNN, respectively, corresponding to improvements of 22.78% and 42.85% relative to ProteinMPNN and HighMPNN. Median pLDDT values were also higher for CyclicMPNN designed sequences (77.51 vs. 75.12 and 73.03, respectively). Across all backbone generation methods and peptide lengths tested (6, 8, 10, and 14 residues), CyclicMPNN consistently showed a higher likelihood of producing sequences compatible with the target backbone structure.

### Sequence generation when starting from specified motifs

Motif inpainting is a common *de novo* design strategy in which a functional segment of a peptide, small molecule, or protein is preserved and used as a structural constraint while generating a surrounding scaffold. The scaffold can adopt diverse conformations, provided that the motif retains its geometry. To evaluate CyclicMPNN’s ability to assign sequences to scaffolds while maintaining a fixed motif, we selected a loop segment from Nrf2, which is known to bind and inhibit Keap1 and has previously been used as a template for cyclic peptide binder design [17, 23].

We based our motif off of the hot loop from Gavenonis et al. [30] with sequence DEETGE from Nrf2, but trimmed it down to EETG, similarly as Rettie et al. 2024 (Fig. 2C; highlights our workflow). We sampled 100 eight-residue cyclic peptide scaffolds containing this motif using the homochiral sampling option available in CyclicCAE (publication in progress) [31]. Sequences were designed for the same set of scaffolds using both CyclicMPNN and ProteinMPNN. Two datasets were generated: one in which only non-motif residues were designed (conditional generation), and another in which the full peptide sequence was redesigned (unconditional generation). This allowed comparison of performance under both constrained and unconstrained sequence design settings.

Results for both methods and their corresponding P_*Near*_ values are shown in Supplementary Fig. S2. In both the full-sequence and non-motif design tasks, CyclicMPNN generated sequences sampled higher P_*Near*_ values overall. Notably, nearly all outlier samples with P_*Near*_ *≥* 0.6 originated from CyclicMPNN designs, with seven such sequences compared to one from ProteinMPNN; in that case, the full sequence required redesign. Examples of high-P_*Near*_ sequences are shown in Fig. 2D, with their corresponding folding funnels in Fig. 2E.

Among the 100 generated sequences, CyclicMPNN produced twice as many stable peptides as ProteinMPNN based on P_*Near*_ funnel analysis (12 vs. 6; Fig. 2F). When comparing sequence uniqueness, ProteinMPNN generated a larger number of unique sequences overall. However, ProteinMPNN also produced more repeated sequences, with some individual sequences occurring 6–8 times (sequences and sequence counts found in Supplementary Table S1 & S2).

### Sequence Space Exploration

Sequence space exploration was performed for sequences generated by both CyclicMPNN and ProteinMPNN. Test-set backbones of varying lengths (6, 8, and 10 amino acids) were selected and used as inputs for sequence design. One sequence was generated per backbone, and the resulting sequences were embedded using Evolutionary Scale Modeling (ESM) to obtain amino acid token representations. A UMAP dimensionality reduction was applied to these embeddings for visualization (Supplementary Fig. S3).

For 6- and 8-mers, CyclicMPNN-generated sequences occupied a broader region of the projected embedding space compared to ProteinMPNN sequences. In contrast, for 10-mers, both methods sampled similar regions of sequence space. Additionally, sequences designed for PDB backbones 9CDT, 9HGC, 9CDU, and 9HGD were included in the analysis. For these structures, CyclicMPNN and ProteinMPNN sequences largely occupied overlapping regions of the embedding space, whereas HighMPNN sequences sampled more distinct regions.

Although CyclicMPNN sequence distributions overlapped with those from ProteinMPNN, CyclicMPNN also sampled adjacent regions not extensively populated by ProteinMPNN. This suggests that, for a given set of backbones, CyclicMPNN explores alternative areas of sequence space that remain compatible with the target structures.

## Methods

### Post-Process & Root Mean Square Deviation Measurement

A consistent sequence design workflow was used throughout this study. For each backbone, five sequences were generated using the respective MPNN model. The sequence with the best model score among the five was selected and folded using HighFold for monomeric peptides and HighFold Multimer for binders. HighFold produced five predicted structures along with associated confidence scores. Backbone RMSD values were calculated by measuring distances between backbone heavy atoms of the predicted structures and those of the original backbone. For each design, the lowest RMSD among the HighFold predictions was used for analysis.

### Generalized Kinematic loop Closure

Generalized Kinematic loop Closure (GenKIC) is a state-of-the-art method for generating cyclic peptide poly-glycine backbones *in silico*. This method is part of the Rosetta macromolecular modeling suite and is based on a robotics-inspired kinematic approach adapted for loop conformation sampling [32, 33]. We implemented a custom PyRosetta script to generate backbone ensembles, which will be made available in the repository described in the Data Availability section.

The number of sampled structures (nstruct) varied by peptide length (see Dataset Creation). Backbone torsions were randomized by allowing the FoldTree root to sample using the `RandomizeBBByRamaPrePro` mover. Ramachandran sampling was performed using the `randomize_backbone_by_rama_prepro` lookup table, followed by filtering with `rama_prepro_check` using a Ramachandran energy cutoff of 2. Backbone–backbone hydrogen bond filters were applied based on peptide length (see Dataset Creation).

### Dataset Creation

Cyclic poly-alanine peptides of lengths 6, 8, and 10 residues were generated using GenKIC. A total of 20,000, 50,000, and 100,000 backbones were sampled for 6-, 8-, and 10-mers, respectively. A backbone–backbone hydrogen bond filter was applied, requiring at least 1, 2, and 3 hydrogen bonds for 6-, 8-, and 10-mers, respectively. After filtering, 19,994, 28,671, and 38,122 conformations remained.

ProteinMPNN was used to design sequences for the sampled poly-alanine structures. For each structure, five sequences were generated, and the sequence with the best model score was selected for folding. HighFold was used to predict structures for all sequences. Sequences with backbone RMSD*≤* 0.35 Å for 6- and 8-mers, or *≤* 0.5 Å for 10-mers, relative to their input backbones were included in the fine-tuning dataset. Additionally, 80 cyclic peptide structures from the HighMPNN training dataset, which were from the PDB, were incorporated into the fine-tuning set.

### Fine-Tuning ProteinMPNN

The fine-tuning dataset, consisting of both *in silico* generated and experimentally determined cyclic peptide structures in PDB format, that were converted into PyTorch tensors. Sequences were clustered using MMseqs2 at 0.3 sequence identity through the HyperMPNN pipeline to group similar sequences and reduce over representation during training. The combined dataset was used to fine-tune ProteinMPNN, with default parameters (Supplementary Table S3), for 200 additional epochs starting from the original pretrained weights. The model checkpoint with the lowest perplexity, epoch 164, during fine-tuning was selected for inference.

### *De Novo* Design & Assessment

For *de novo* evaluation, 10,000 cyclic poly-alanine backbones of lengths 6, 8, and 10 residues were generated using GenKIC and filtered to retain structures with at least 1, 2, and 3 backbone hydrogen bonds for 6-, 8-, and 10-mers, respectively. ProteinMPNN and CyclicMPNN each generated five sequences per backbone, and the sequence with the best model score was selected. HighFold produced five predicted structures per sequence. RMSD was calculated between the backbone heavy atoms of the predicted structures and the corresponding input poly-alanine backbone.

For larger peptides, RFPeptide was used to generate 100 cyclic 14-mer poly-glycine backbones. With arguments `inference.cyclic=True, contigmap.contigs=[14-14]`. ProteinMPNN, HighMPNN, and CyclicMPNN each generated five sequences per backbone. The sequence with the best model score was folded using HighFold, and RMSD was measured between the predicted structure and the original poly-glycine backbone.

## Discussion

In this work, we present CyclicMPNN, a fine-tuned variant of ProteinMPNN developed specifically for the design of L-*α*-amino acid cyclic peptide sequences. ProteinMPNN was originally trained on X-ray structures from the PDB above a minimum residue length threshold, which results in limited representation of small cyclic peptides. This under representation makes cyclic peptide sequence design a challenging task for the original model. Here, we demonstrate that datasets generated entirely through *in silico* methods can be used to meaningfully extend the applicability of pretrained protein design models.

Despite being trained predominantly on *in silico* generated structures, CyclicMPNN accurately recapitulates experimentally determined X-ray crystal structures, producing sequences that fold into conformations closely matching the native backbones. In addition to recovering known structural motifs, CyclicMPNN samples sequences that are distinct from previously observed sequences for the same structures, indicating exploration of alternative compatible regions of sequence space.

Because ProteinMPNN is widely used in *de novo* sequence design settings, we directly compared performance in this domain. Across cyclic peptides of 6, 8, 10, and 14 residues, CyclicMPNN consistently outperformed ProteinMPNN in structural reconstruction accuracy (RMSD) and model confidence (pLDDT). For 6-mers, CyclicMPNN-designed sequences also exhibited improved stability metrics, as measured by P_*Near*_. Furthermore, CyclicMPNN showed improved performance in motif-constrained (inpainting) sequence design tasks.

Overall, this work demonstrates that cyclic peptide design methods can be substantially improved using carefully constructed *in silico* datasets. More broadly, our fine-tuning framework suggests a general strategy for adapting large pretrained protein design models to under represented design tasks, potentially enabling the development of new specialized design tools for other classes of constrained biomolecular systems.

## Author contributions

ACP conceived of the presented idea. ACP developed the theory while ACP and YJ performed the computations. ACP and PH verified the analytical methods. PH provided mentorship for both ACP and YJ. ACP and YJ made the figures for the manuscript. ACP and YJ made the original first draft, while all authors provided editing and feedback.

## Data Availability

All weights and scripts required to recreate the CyclicMPNN GenKIC training dataset are available at ParisaH-Lab/CyclicMPNN.

## Acknowledgments

ACP and YJ thank the University of Oregon, as all of the the computations reported in this paper were performed using resources made available by the University of Oregon. In particular the HPC Cluster computational resources, Talapas, at the University of Oregon. ACP thanks the scientists P. Doug Renfrew and Vikram K. Mulligan at the Flatiron Institute, for their helpful conversations and feedback. PH, ACP, and YJ are funded through the NIH grant DP2GM146249f.

