## Supplemental Information for "CyclicMPNN: Stable Cyclic Peptide Sequence Generation"

### Supplementary Information

#### $P_{Near}$ Stability Test

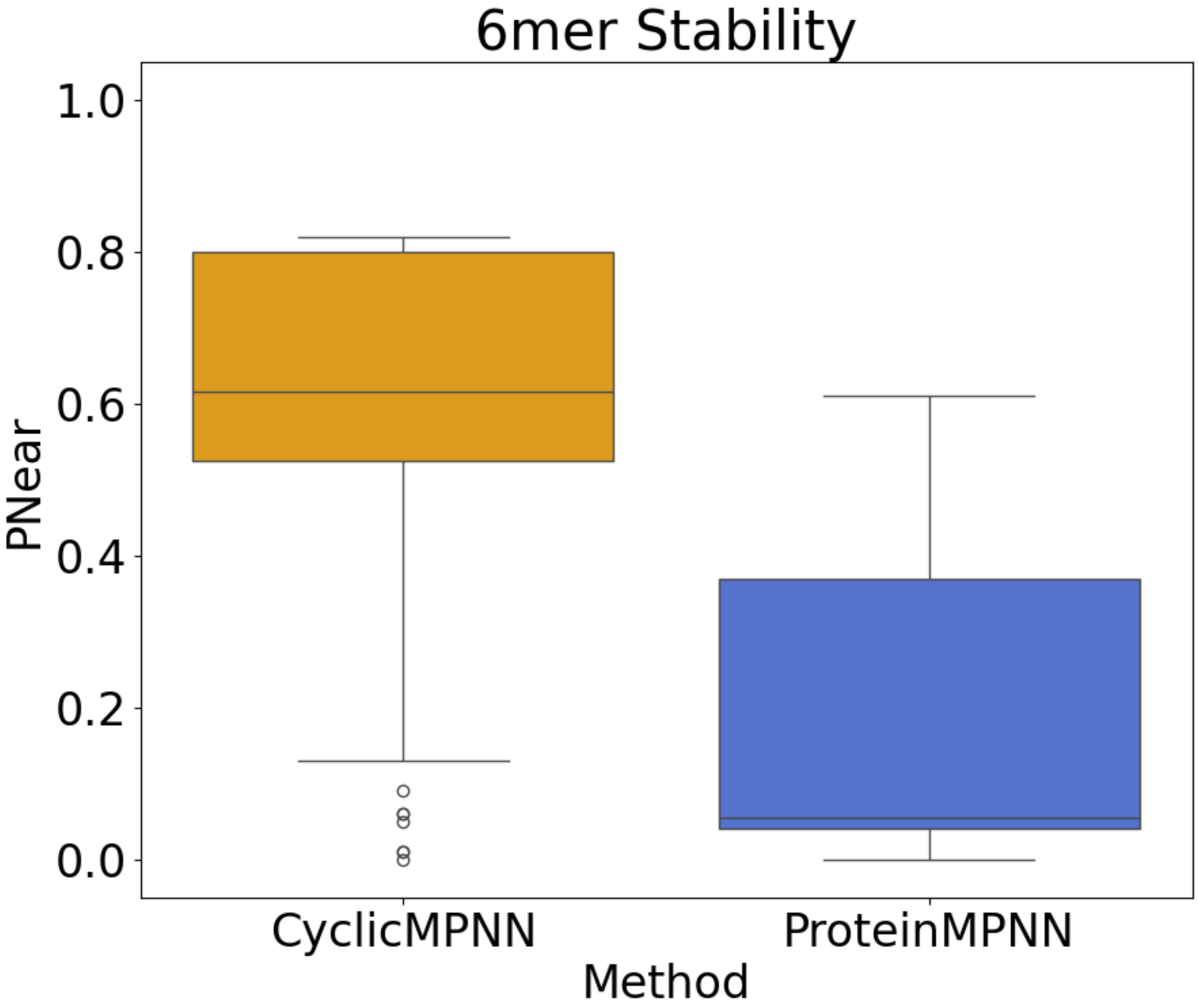


**Figure S1:** 6mer PN ear values obtained using simple_cycpep_predict. It can be seen that sequences from CyclicMPNN have a much higher proportion of stable structures than sequences ProteinMPNN produced.

Nrf2 $P_{Near}$ Comparison of Full Sequence vs. Non-Motif Sequence

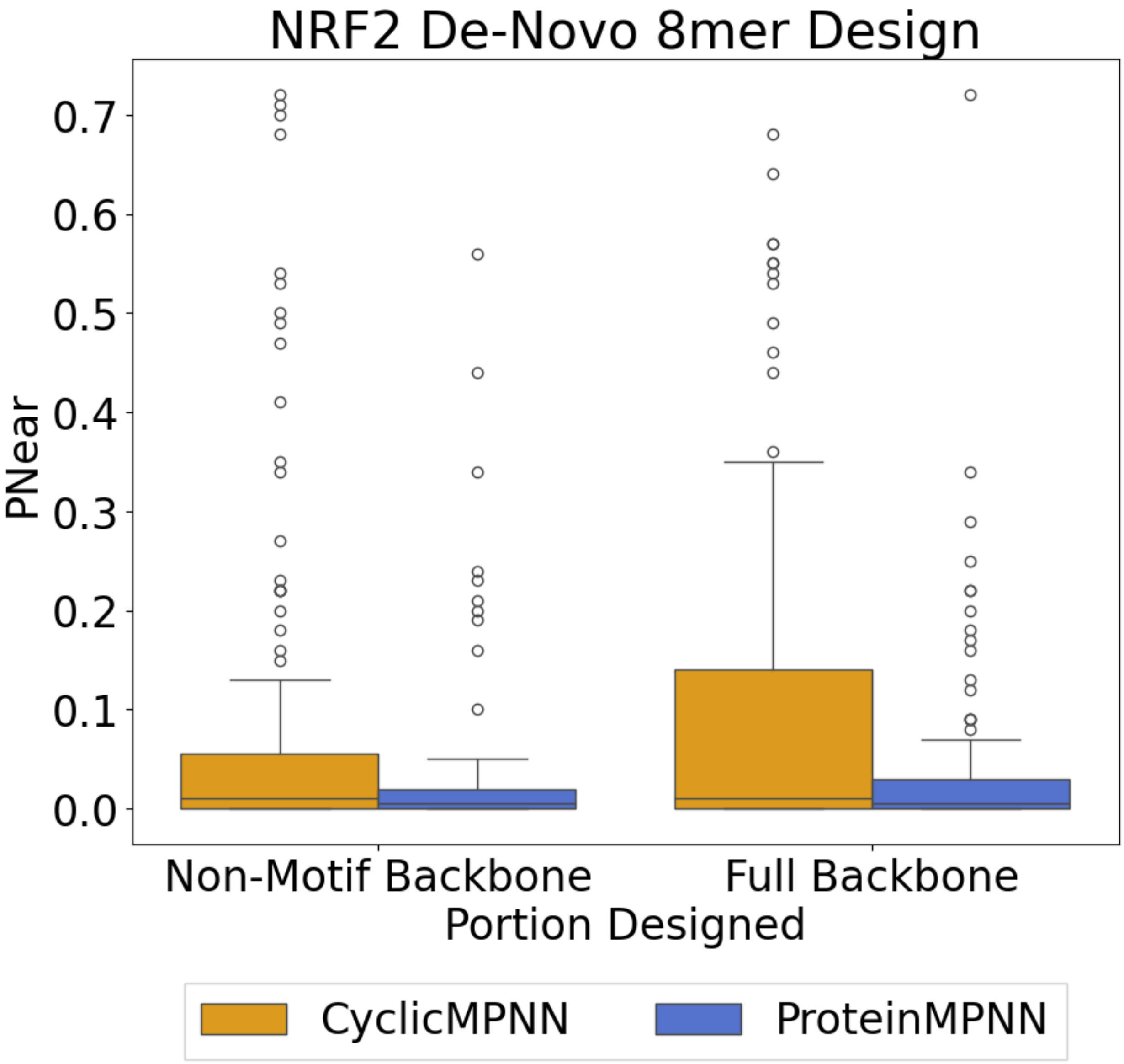


**Figure S2**: Motif-inpainting of the Nrf2 loop that inhibits Keap1 using CyclicMPNN and proteinMPNN sequence stability are shown here. This shows sequence design for non-motif portions of the cyclic scaffolds on the left, and the right shows full backbone sequence redesign. It can be seen that a larger portion of outliers are ≥ 0.6.

### Nrf2 Non-Motif Inpainting Sequence Counts

ProteinMPNN and CyclicMPNN sequences were extracted and counted for redundancy. Below are tables of ProteinMPNN and CyclicMPNN unique sequences and the amount of times those sequences show up.

**Table S1**: ProteinMPNN Sequences

| **Sequence** | **Count** |
|---|---|
| HIS-GLU-GLU-THR-GLY-THR-GLY-PRO | 2 |
| TYR-GLU-GLU-THR-GLY-GLY-GLU-LYS | 2 |
| ILE-GLU-GLU-THR-GLY-ASN-LEU-PRO | 1 |
| ASP-GLU-GLU-THR-GLY-GLY-GLU-ARG | 2 |
| VAL-GLU-GLU-THR-GLY-GLY-GLU-LYS | 7 |
| ASN-GLU-GLU-THR-GLY-THR-ALA-PRO | 1 |
| ILE-GLU-GLU-THR-GLY-GLU-LYS-GLU | 1 |
| TYR-GLU-GLU-THR-GLY-THR-GLY-PRO | 1 |
| VAL-GLU-GLU-THR-GLY-GLY-GLU-ARG | 1 |
| ASN-GLU-GLU-THR-GLY-ASP-PRO-ARG | 1 |
| VAL-GLU-GLU-THR-GLY-HIS-PRO-LEU | 1 |
| GLY-GLU-GLU-THR-GLY-THR-GLY-GLU | 1 |
| LEU-GLU-GLU-THR-GLY-HIS-GLU-LYS | 1 |
| ILE-GLU-GLU-THR-GLY-ASP-GLN-PRO | 1 |
| ARG-GLU-GLU-THR-GLY-GLU-GLU-GLU | 1 |
| ILE-GLU-GLU-THR-GLY-ASP-LEU-PRO | 1 |
| ASN-GLU-GLU-THR-GLY-ILE-GLY-LYS | 1 |
| SER-GLU-GLU-THR-GLY-GLU-GLY-PRO | 1 |
| ILE-GLU-GLU-THR-GLY-HIS-PRO-LEU | 1 |
| ASN-GLU-GLU-THR-GLY-THR-PRO-GLU | 6 |
| VAL-GLU-GLU-THR-GLY-SER-THR-THR | 1 |
| ILE-GLU-GLU-THR-GLY-THR-LYS-PRO | 1 |
| ILE-GLU-GLU-THR-GLY-HIS-PRO-GLU | 1 |
| GLU-GLU-GLU-THR-GLY-GLY-GLU-LYS | 8 |
| VAL-GLU-GLU-THR-GLY-GLY-GLU-GLU | 1 |
| ARG-GLU-GLU-THR-GLY-ASN-GLY-PRO | 1 |
| ILE-GLU-GLU-THR-GLY-ASN-VAL-PRO | 2 |
| ASN-GLU-GLU-THR-GLY-THR-GLY-VAL | 2 |

| | |
|---|---|
| ASN-GLU-GLU-THR-GLY-GLY-GLU-LYS | 4 |
| ASN-GLU-GLU-THR-GLY-ASN-LEU-PRO | 2 |
| ILE-GLU-GLU-THR-GLY-THR-ALA-LYS | 1 |
| VAL-GLU-GLU-THR-GLY-GLU-LYS-GLU | 1 |
| LEU-GLU-GLU-THR-GLY-HIS-PRO-GLU | 2 |
| ASN-GLU-GLU-THR-GLY-ASN-GLN-ALA | 2 |
| VAL-GLU-GLU-THR-GLY-GLU-GLY-GLU | 1 |
| LEU-GLU-GLU-THR-GLY-THR-GLY-PRO | 1 |
| ARG-GLU-GLU-THR-GLY-GLU-LYS-GLU | 1 |
| ARG-GLU-GLU-THR-GLY-HIS-PRO-GLU | 2 |
| ASN-GLU-GLU-THR-GLY-GLY-GLU-GLU | 3 |
| THR-GLU-GLU-THR-GLY-GLU-ALA-GLY | 1 |
| ILE-GLU-GLU-THR-GLY-GLN-LYS-GLU | 1 |
| SER-GLU-GLU-THR-GLY-THR-GLY-GLY | 1 |
| ILE-GLU-GLU-THR-GLY-SER-LYS-GLU | 1 |
| ASN-GLU-GLU-THR-GLY-THR-ALA-GLU | 1 |
| ILE-GLU-GLU-THR-GLY-SER-GLU-LYS | 1 |
| SER-GLU-GLU-THR-GLY-GLU-GLY-ARG | 1 |
| ASN-GLU-GLU-THR-GLY-THR-LYS-GLU | 2 |
| ASN-GLU-GLU-THR-GLY-THR-LYS-PRO | 1 |
| VAL-GLU-GLU-THR-GLY-THR-GLY-PRO | 2 |
| ALA-GLU-GLU-THR-GLY-ASP-PRO-ARG | 1 |
| ASN-GLU-GLU-THR-GLY-GLU-GLY-GLU | 1 |
| ARG-GLU-GLU-THR-GLY-THR-GLY-GLY | 1 |
| ARG-GLU-GLU-THR-GLY-THR-GLU-GLU | 1 |
| ARG-GLU-GLU-THR-GLY-GLU-GLU-THR | 1 |
| THR-GLU-GLU-THR-GLY-GLY-GLU-LYS | 2 |
| ASN-GLU-GLU-THR-GLY-HIS-PRO-LEU | 1 |
| VAL-GLU-GLU-THR-GLY-GLY-ALA-GLU | 1 |
| ILE-GLU-GLU-THR-GLY-THR-GLY-PRO | 1 |
| ASN-GLU-GLU-THR-GLY-ASN-GLY-VAL | 1 |
| ARG-GLU-GLU-THR-GLY-ILE-PRO-GLU | 1 |
| ILE-GLU-GLU-THR-GLY-LEU-PRO-GLU | 1 |
| ILE-GLU-GLU-THR-GLY-THR-GLN-PRO | 1 |
| SER-GLU-GLU-THR-GLY-GLU-GLY-GLY | 1 |

| | |
|---|---|
| ILE-GLU-GLU-THR-GLY-ASN-GLN-PRO | 1 |
| ILE-GLU-GLU-THR-GLY-GLY-GLU-LYS | 1 |

**Table S2**: CyclicMPNN Sequences

| Sequence | Count |
|---|---|
| SER-GLU-GLU-THR-GLY-ASP-GLY-PRO | 1 |
| ASN-GLU-GLU-THR-GLY-SER-GLU-ILE | 2 |
| ASN-GLU-GLU-THR-GLY-GLY-LEU-PRO | 2 |
| ASN-GLU-GLU-THR-GLY-LEU-TYR-ASP | 1 |
| ASN-GLU-GLU-THR-GLY-ASP-TYR-ILE | 3 |
| ILE-GLU-GLU-THR-GLY-ILE-CYS-PRO | 1 |
| ILE-GLU-GLU-THR-GLY-LEU-VAL-GLU | 1 |
| ILE-GLU-GLU-THR-GLY-ILE-TYR-PRO | 2 |
| ILE-GLU-GLU-THR-GLY-HIS-PRO-ILE | 4 |
| ASN-GLU-GLU-THR-GLY-HIS-PRO-GLU | 4 |
| HIS-GLU-GLU-THR-GLY-ASN-GLY-PRO | 1 |
| ILE-GLU-GLU-THR-GLY-HIS-PRO-GLU | 2 |
| GLY-GLU-GLU-THR-GLY-ASP-GLY-ASP | 1 |
| ILE-GLU-GLU-THR-GLY-LEU-PRO-ASP | 1 |
| ASN-GLU-GLU-THR-GLY-ASP-LEU-PRO | 2 |
| TYR-GLU-GLU-THR-GLY-HIS-PRO-ILE | 2 |
| ASN-GLU-GLU-THR-GLY-PRO-LEU-PRO | 1 |
| ASN-GLU-GLU-THR-GLY-ILE-GLY-GLY | 1 |
| PHE-GLU-GLU-THR-GLY-ILE-GLY-PRO | 1 |
| ASN-GLU-GLU-THR-GLY-GLY-PHE-ILE | 3 |
| ILE-GLU-GLU-THR-GLY-HIS-PRO-LEU | 2 |
| ASN-GLU-GLU-THR-GLY-THR-PRO-GLU | 2 |
| TYR-GLU-GLU-THR-GLY-SER-LYS-ASP | 1 |
| ASN-GLU-GLU-THR-GLY-THR-TYR-PRO | 1 |
| ASN-GLU-GLU-THR-GLY-SER-TYR-GLU | 1 |
| TYR-GLU-GLU-THR-GLY-LEU-PRO-GLU | 2 |
| HIS-GLU-GLU-THR-GLY-ASN-LEU-PRO | 5 |
| ASN-GLU-GLU-THR-GLY-THR-GLY-ASP | 1 |
| ASN-GLU-GLU-THR-GLY-GLY-VAL-ASP | 3 |
| ASN-GLU-GLU-THR-GLY-ASN-LEU-PRO | 1 |

| | |
|---|---|
| ASP-GLU-GLU-THR-GLY-HIS-PRO-ILE | 5 |
| ILE-GLU-GLU-THR-GLY-ILE-ALA-ASP | 1 |
| ILE-GLU-GLU-THR-GLY-LEU-PRO-GLU | 3 |
| HIS-GLU-GLU-THR-GLY-PRO-GLY-ASP | 1 |
| HIS-GLU-GLU-THR-GLY-ASP-GLY-PRO | 2 |
| ILE-GLU-GLU-THR-GLY-ILE-VAL-GLY | 1 |
| ARG-GLU-GLU-THR-GLY-LEU-PRO-GLU | 1 |
| THR-GLU-GLU-THR-GLY-ASP-ILE-PRO | 1 |
| ILE-GLU-GLU-THR-GLY-LEU-ARG-PRO | 2 |
| TYR-GLU-GLU-THR-GLY-ASN-LEU-GLY | 1 |
| ILE-GLU-GLU-THR-GLY-SER-VAL-ILE | 2 |
| SER-GLU-GLU-THR-GLY-SER-PRO-ILE | 1 |
| ASN-GLU-GLU-THR-GLY-LEU-PRO-GLU | 2 |
| THR-GLU-GLU-THR-GLY-LEU-GLY-ASN | 1 |
| ARG-GLU-GLU-THR-GLY-ASP-TYR-ILE | 2 |
| ILE-GLU-GLU-THR-GLY-HIS-PRO-ASP | 1 |
| SER-GLU-GLU-THR-GLY-LEU-TYR-PRO | 1 |
| ASN-GLU-GLU-THR-GLY-HIS-PRO-ILE | 2 |
| ASP-GLU-GLU-THR-GLY-THR-GLY-ASP | 1 |
| ASN-GLU-GLU-THR-GLY-GLY-VAL-ILE | 2 |
| ASN-GLU-GLU-THR-GLY-LEU-GLY-ASP | 1 |
| ASN-GLU-GLU-THR-GLY-GLY-VAL-GLU | 1 |
| TYR-GLU-GLU-THR-GLY-ASN-GLY-PRO | 1 |
| THR-GLU-GLU-THR-GLY-THR-GLY-ARG | 1 |
| SER-GLU-GLU-THR-GLY-SER-GLU-ILE | 1 |
| ILE-GLU-GLU-THR-GLY-LEU-TYR-PRO | 1 |
| ASN-GLU-GLU-THR-GLY-LEU-GLY-VAL | 1 |
| ASN-GLU-GLU-THR-GLY-GLY-TYR-ASP | 1 |
| LEU-GLU-GLU-THR-GLY-HIS-PRO-GLU | 1 |
| ILE-GLU-GLU-THR-GLY-HIS-PRO-VAL | 1 |
| GLY-GLU-GLU-THR-GLY-ASP-GLY-GLY | 1 |
| TYR-GLU-GLU-THR-GLY-GLY-LEU-PRO | 1 |

### Latent Space Exploration

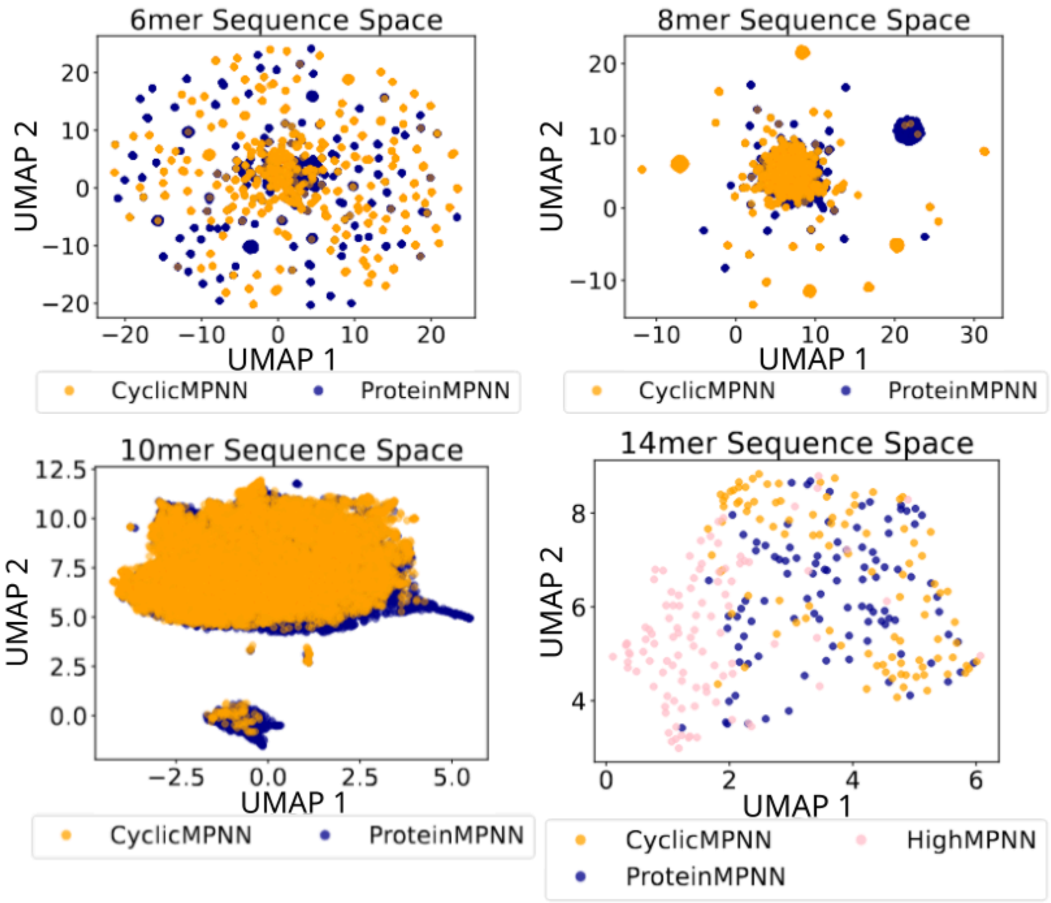


**Figure S3:** Sequences generated for all GenKIC created backbones (6mer, 8mer, and 10mers) and RFPeptide generated structures (14mer) sequences were extracted and run through ESM to grab their latent embedding. Then scikit-learn's UMAP function was used to embed them into a 2 dimension latent space. Here we can seeCyclicMPNN, ProteinMPNN, and HighMPNN sequence latent-space representations in orange, blue, and pink, respectively.

### Alternating the number of neighbors when fine-tuning ProteinMPNN

| # of Neighbors | Epoch | Train Perplexity | Validation Perplexity | Train Accuracy | Validation Accuracy |
|---|---|---|---|---|---|
| 3 | 238 | 2.668 | 6.504 | 0.697 | 0.461 |
| 4 | 223 | 2.283 | 5.245 | 0.748 | 0.534 |
| **48** | **164** | **2.151** | **3.91** | **0.772** | **0.566** |

**Table S3:** We tried varying sizes of neighbors to include while fine-tuning ProteinMPNN, we checked to see if reducing the number of neighbors would be beneficial for the smaller cyclic peptides. However, it seemed that using default, 48, showed the best across the border metrics in total. Though in particular here we are showing the lowest validation perplexity of the 200 epochs. The row in bold are the weights used for CyclicMPNN.
